# Protein evolution is structure dependent and non-homogeneous across the tree of life

**DOI:** 10.1101/2020.01.28.923458

**Authors:** Akanksha Pandey, Edward L. Braun

## Abstract

**Motivation:** Protein sequence evolution is a complex process that varies among-sites within proteins and across the tree of life. Comparisons of evolutionary rate matrices for specific taxa (‘clade-specific models’) have the potential to reveal this variation and provide information about the underlying reasons for those changes. To study changes in patterns of protein sequence evolution we estimated and compared clade-specific models in a way that acknowledged variation within proteins due to structure.

**Results:** Clade-specific model fit was able to correctly classify proteins from four specific groups (vertebrates, plants, oomycetes, and yeasts) more than 70% of the time. This was true whether we used mixture models that incorporate relative solvent accessibility or simple models that treat sites as homogeneous. Thus, protein evolution is non-homogeneous over the tree of life. However, a small number of dimensions could explain the differences among models (for mixture models ~50% of the variance reflected relative solvent accessibility and ~25% reflected clade). Relaxed purifying selection in taxa with lower long-term effective population sizes appears to explain much of the among clade variance. Relaxed selection on solvent-exposed sites was correlated with changes in amino acid side-chain volume; other differences among models were more complex. Beyond the information they reveal about protein evolution, our clade-specific models also represent tools for phylogenomic inference.

**Availability:** Model files are available from https://github.com/ebraun68/clade_specific_prot_models.

**Supplementary information:** Supplementary data are appended to this preprint.

## 1 Introduction

In phylogenetics, tree topology and/or branch lengths are typically the parameters of interest. However, rate matrices estimated as part of maximum likelihood (ML) analyses also provide information about the process of evolution. Focusing on protein evolution, patterns of substitution processes vary across the tree of life and among proteins (e.g., Braun, 2018; Weber and Whelan, 2019; Zou and Zhang, 2019). It has long been appreciated (Kimura, 1986) that the accumulation of substitutions over evolutionary time reflects two processes: 1) the rate at which novel mutations enter populations; and 2) the impact of drift and selection on the fate of those mutations. This paradigm suggests the patterns of protein evolution will vary across the tree of life; after all, the rate and spectrum of mutations and strength of selection (the latter reflecting, in large part, variation in effective population size, *N*_e_) varies across the tree (Sung *et al*., 2012; Behringer and Hall, 2016). The sensitivity of ratio of radical to conservative amino acid substitutions to *N*_e_ (Nabholz *et al*., 2013; Weber and Whelan, 2019) suggests variation in the strength of selection is likely to be especially important for establishing the patterns of protein evolution.

Using the radical to conservative substitution ratio to examine changes in the pattern of sequence evolution is complicated by the challenge of defining radical amino acid changes. Even in the earliest days of molecular evolution Zuckerkandl and Pauling (1965, p. 129) recognized that the “…inadequacy of *a priori* views on [amino acid substitution] conservatism and nonconservatism is patent”; that problem remains unsolved. Many studies divide residues into two categories (e.g., polar/non-polar or small/large) and treat between-category substitutions as radical (e.g., Nabholz *et al*., 2013). That idea can be extended by using continuous values to describe the physicochemical characteristics of the amino acids instead of binary classification (Braun, 2018), but that still relies on the use of prespecified amino acid characteristics. Assessing changes in the process of protein sequence evolution without *a priori* assumptions would be desirable. In principle, one could use the general Markov model (GMM) to estimating amino acid rate matrices. However, using the amino acid GMM requires estimation of 380 free parameters per branch, unlike the nucleotide GMM which only requires 12 free parameters (Barry and Hartigan, 1987). Moreover, the GMM cannot be used with among-sites rate variation (except for a +invariant sites version; Jayaswal *et al*., 2007) and rate variation is ubiquitous in protein evolution (Echave *et al*., 2006). Thus, the GMM cannot be used for this purpose.

The general time-reversible model for amino acids (GTR_20_) might provide a feasible framework for parameter estimation. The GTR_20_ instantaneous rate matrix (usually called the ***Q*** matrix) can be decomposed into a symmetric rate (***R***) matrix with 189 free parameters that reflect the ‘exchangeability’ of each pair of amino acids and a diagonal matrix (**Π**) with 19 free parameters for equilibrium amino acid frequencies (Le and Gascuel, 2008). Of course, the time-reversibility assumption that limits the number of free parameters is inappropriate when protein evolution has changed across the tree; after all, postulating that the model changes over time (i.e., that the model is non-homogeneous) intrinsically renders models non-time-reversible. However, we can avoid this problem by estimating GTR_20_ parameters for clades with a limited taxonomic scope and then comparing the clade-specific parameter estimates. If the deviations from time-reversibility for the underlying model of protein evolution are sufficiently limited within clades comparisons among those clades should reveal the ways protein evolution has changed across the tree of life.

Using GTR_20_ parameter estimates to understand shifts in process of sequence evolution presents several challenges. Although previous studies (Huzurbazar *et al*., 2010; Zou and Zhang, 2019) indicate that we will find differences among clades, the complex and heterogeneous nature of protein evolution suggests there will also be substantial variation among proteins (Braun, 2018) and among sites within proteins (Meyer and Wilke, 2013; Echave *et al*., 2016). There are two way this heterogeneity could confound our ability the use of GTR_20_ parameter estimates to understand patterns of protein evolution across the tree of life. First, a high degree of variation among individual proteins might obscure variation among clades (Fig. 1). Second, simply optimizing GTR_20_ model parameters on a large protein dataset will yield average exchangeability estimates for all sites. If there is substantial variation among-sites within proteins the patterns revealed by comparing these ‘averaged’ parameter estimates could be confusing. These factors make it important to find ways to examine the impact of these sources of variation on any conclusions we reach regarding differences among taxa.

**Fig. 1.**
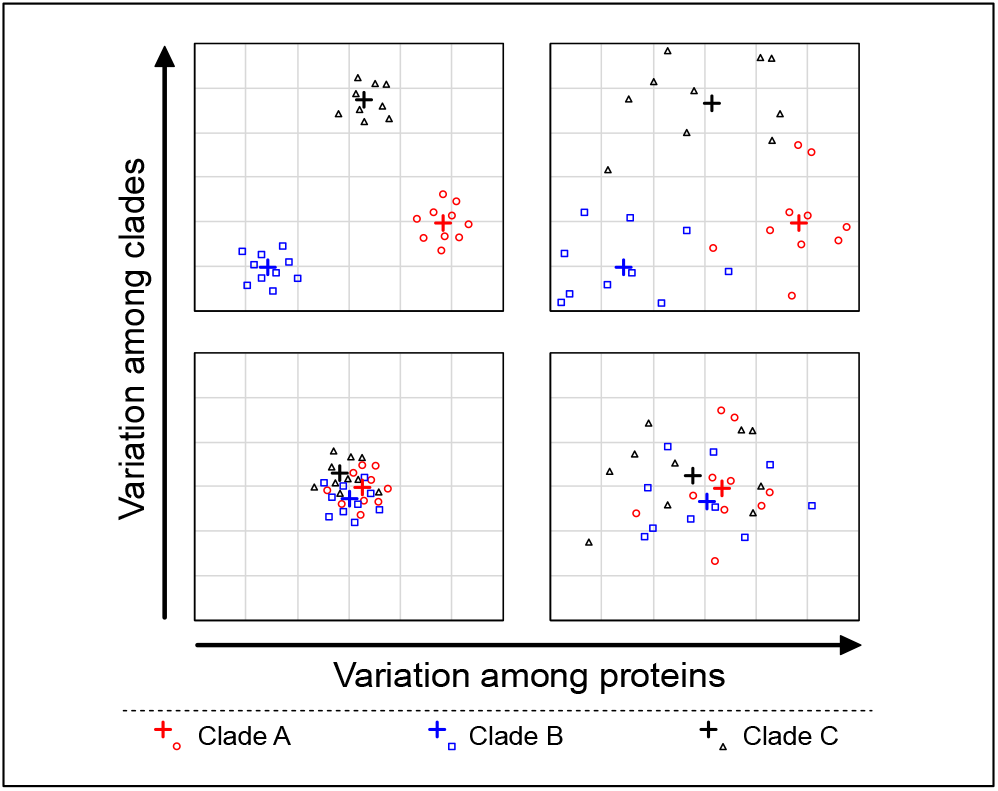
Possible patterns for the variation in patterns of protein sequence evolution, both among proteins and among clades. Conceptual illustration showing the relationships among proteins in the underlying models of presented after some type of dimension reduction. The crosses indicate models generated using the training data (i.e., large-scale averages for the parameters) and the smaller circles, squares, and triangles indicate individual proteins. Obviously, the number of dimensions necessary to provide a good summary of the data in GTR_20_ rate matrices in unclear; we have shown two dimensions in the interest of simplicity.

Assessing fine-scale variation (i.e., variation among individual proteins and among sites within proteins) is challenging. The simple approach of estimating GTR_20_ parameters for individual proteins and comparing them will not work; although the dimension of GTR_20_ is lower than that of the GMM it is still a parameter-rich model and individual proteins will not provide sufficient data for accurate parameter estimates. However, it is possible to estimate model parameters on a relatively large training dataset and then use those parameters to classify proteins in an independent validation dataset. Hereafter, we will call the GTR_20_ matrices estimated as part of this study ‘models’ because they are analogous to the empirical models that are often used in protein phylogenetics, such as the PAM (Dayhoff *et al*., 1978), JTT (Jones *et al*., 1992), WAG (Whelan and Goldman, 2001), and LG (Le and Gascuel, 2008) (we will also call PAM, JTT, WAG, LG, and related models ‘standard empirical models’). Using a combination of clade-specific and standard empirical models to classify individual proteins should make it possible to establish the part of the parameter space shown in Fig. 1 that best describes large-scale patterns of protein evolution. Clade-specific models should fail as a classifier if the variance among individual proteins exceeds the variation among clades (lower portion of Fig. 1). In contrast, if the variation among clades exceeds the variation among proteins (upper portion of Fig 1), we expect the use of models as classifiers to work (i.e., the best-fitting model for validation set proteins will be the model generated from that clade). Finally, the number of times that model fit fails as a classifier will increase as the variation among proteins increases. It should be possible to establish the specific parameters that vary among clades and determine whether they are consistent with predictions regarding the expected differences among clades in the strength of selection, assuming there is sufficient variation among clades.

The other type of fine-scale variation, variation among sites within proteins, is more difficult to examine. Patterns of protein evolution are complex (e.g., Wilke, 2012) and the best way to extract information about the patterns of molecular evolution while still acknowledging variation within proteins remains unclear. However, selection to maintain protein structure, which has a fundamental role in maintaining protein function, is likely to play a major role in establishing the overall patterns within of amino acid substitution matrices (Parisi and Echave, 2001). The relative solvent accessibility (RSA) of individual amino acids is one of the most important determinants of the patterns of sequence evolution for globular proteins (Conant and Stadler, 2009; Pandey and Braun, 2019). This suggest it should be possible to subdivide proteins into solvent exposed (high RSA) and buried (low RSA) sites before estimating substitution matrix parameters for various clades. This would add another dimension to the parameter space shown in Fig. 1 (i.e., a dimension describing variation among sites within proteins). It also makes it necessary to use of a mixture model as a classifier (i.e., a model with two ‘sub-models’ where the site likelihoods are calculated as a weighted mixture of both sub-model matrices). However, using these exposed/buried (‘XB’) mixture models is a straightforward extension of the idea of using models as a classifier to determine which part of parameter space best describe the large-scale patterns of protein evolution.

Herein, we examine the extent to which models of protein sequence evolution exhibit clade-specific features using six eukaryotic datasets selected to exhibit differences in the strength of selection. These clades selected for this study included vertebrates (expected to have small longterm *N*_e_), plants (expected to have intermediate long-term *N*_e_), and microbial eukaryotes (expected to have large long-term *N*_e_). We focused on eukaryote datasets to limit the impact of horizontal gene transfer on our parameter estimates; the high rate of horizontal gene transfer in prokaryotes (Soucy *et al*., 2015) could have distorted estimates. We added a seventh dataset with a broad sample of eukaryotes; multiple changes in the underlying model of sequence evolution are likely to have occurred for the taxa in that dataset (unless the best description of protein evolution actually lies in the lower part of Fig. 1). This ‘all Euk’ dataset was included to assess the impact of using a dataset presumed to have experienced variation on the parameter estimates we obtained. We then used the new models as classifiers to assess among-protein variation and examine the way models differ, examining parameter differences among clades, among sites that were grouped by RSA, and for the combination of RSA and clade.

## 2 Methods

We generated 14 new protein models (seven based on all sites and seven XB mixture models) that were trained using arbitrarily selected proteins from seven published datasets (Table 1). One training dataset (Prum *et al*., 2015) was available as nucleotide sequences, often including only one exon. For that dataset we extracted the coding exons and expanded the dataset by adding orthologous sequences from 117 avian genome assemblies (using the pipeline described in Reddy *et al*., 2017). After adding taxa to the Prum *et al*. (2015) dataset each locus was re-aligned using MAFFT v.7.130b (Katoh *et al*., 2009).

**Table 1.** Training datasets selected for this study

| Clade | # Proteins/Sites | # Taxa | Best Model | Citation |
| --- | --- | --- | --- | --- |
| Birds (1) | 250 / 109,969 | 48 | JTT | Jarvis *et al.* (2015) |
| Birds (2) | 250 / 161,112 | 317 | HIVb | Prum *et al.* (2015) |
| Mammals | 250 / 238,319 | 116 | HIVb | Douzery *et al.* (2014) |
| Plants (1) | 310 / 80,315 | 46 | JTT | Xi *et al.* (2014) |
| Oomycetes | 200 / 81,802 | 17 | LG | Ascunce *et al.* (2017) |
| Yeasts | 277 / 83,312 | 343 | LG | Shen *et al.* (2018) |
| All Euk (1) | 248 / 58,469 | 149 | LG | Strassert *et al.* (2019) |

In all cases, the best-fitting standard empirical model included +F+I+G4. The Prum *et al.* (2015) dataset was modified by adding 117 taxa (see above)

We estimated parameters for the new models using IQ-TREE v. 1.6.10 (Nguyen *et al*., 2015) as implemented in CIPRES science gateway (Miller *et al*. 2010). Before conducting the full model optimization, we identified he best-fitting standard empirical model for each training dataset using the -m TEST option with AIC_*c*_ as the decision criterion. The best-fitting standard empirical model varied among clades (Table 1), but the rate heterogeneity parameters for all best-fit models included both invariant sites and Γ-distributed rates. Thus, we used GTR_20_+I+Γ to estimate the new clade-specific models. We fixed the tree topology and among-sites rate heterogeneity parameters (the Γ-distribution shape parameter and proportion of invariant sites) based on the analysis using the standard empirical model before optimizing fitting GTR_20_+I+Γ model parameters. The new clade-specific models are available from github in a format usable by IQ-TREE and PAML (Yang, 2007).

We selected six validation datasets (Table 2). In most cases, the datasets in Table 1 had enough alignments to divide them into training and validation sets. However, we used all genes in the plant and ‘all Euk’ (the latter includes a broad sample of eukaryotes) datasets. In those two cases, we selected another dataset with a comparable set of taxa to use as the validation set. We used BLAST (Camacho *et al*., 2009) to search the validation dataset with training set queries and we removed proteins with a low *E*-value (our cut-off was 10^−40^). This eliminated any overlap between these two training and validation datasets. We determined the best-fitting model for each protein in the validation sets by using the -mset option to supply a list of all 18 standard empirical models as well as the seven new models estimated for this study to IQ-TREE. For comparison, we conducted the same analyses using the proteins in training datasets.

**Table 2.**
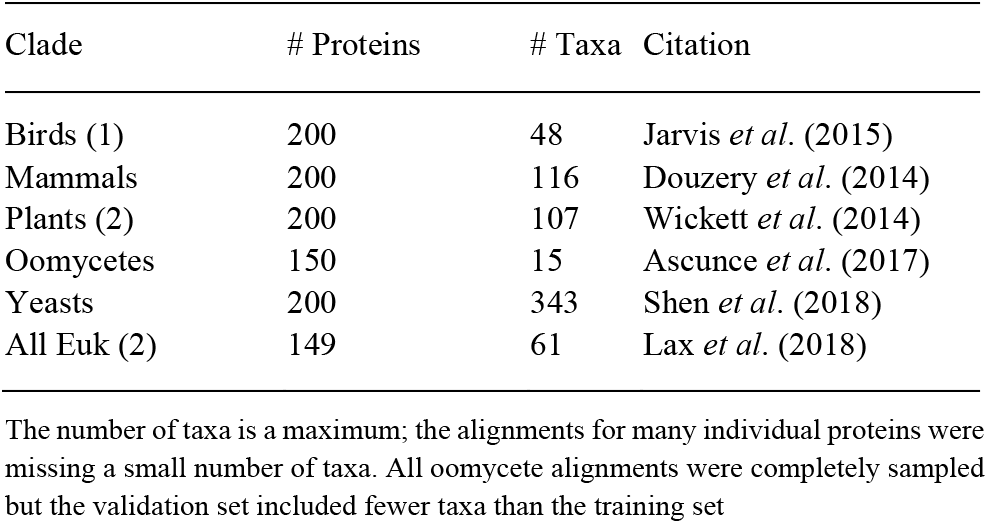
Validation datasets selected for this study

To reduce within-protein heterogeneity, we separated globular proteins into exposed and buried sites. First, we used TMHMM (Krogh *et al*., 2001) to identify and remove transmembrane proteins. Then the globular proteins were subdivided based on site RSA using a pipeline available from https://github.com/aakanksha12/Structural_class_assignment_pipeline. The pipeline generates a weighted consensus sequence that is then used as input for ACCpro (Pollastri *et al*., 2002) from the SCRATCH-1D suite (Magnan and Baldi, 2014). ACCpro assigns each residue to one of the two categories: exposed or buried, with the latter defined as <25% RSA. This information is written to a nexus file (Maddison *et al*., 1997) for each protein multiple sequence alignment that includes charsets for the two RSA classes (solvent exposed and buried). Finally, PAUP* 4.0b10 (Swofford, 2003) was used extract the data subsets defined by RSA. Once the data were subdivided into exposed and buried residues the data for all proteins were concatenated, resulting in 14 data matrices (one exposed dataset and one buried dataset for the seven datasets in Table 1).

Rate matrices were estimated for all 14 exposed- and buried-site training datasets as described above for the all sites datasets. The exposed and buried rate matrices were then combined to create a set of seven XB mixture models, in which the site likelihoods are weighted averages over both alternative (exposed and buried) matrices. We identified the best-fitting mixture model for individual proteins in the validation datasets using ‘fit_mixture_model.pl’ (available from github), which examines the fit of eight models: the seven XB models we generated and the Le *et al*. (2008) EX2 model (which is also an exposed/buried mixture model). We tested four versions of each model that differed in their treatment of rate heterogeneity (no rate heterogeneity vs. Γ-distributed rates) and equilibrium amino acid frequencies (the used of matrix frequencies vs optimized [+FO] amino acid frequencies). As above, we also assessed model fit for all proteins in training datasets. The XB models are available from github as a nexus file that can be used by IQ-TREE.

We analyzed the GTR_20_ exchangeability (***R***) matrices for standard empirical models and the new models generated in two ways. First, we clustered the distances among matrices by neighbor-joining (Saitou and Nei, 1987). Second, we used principal component analysis (PCA) to explore differences among models. We normalized the matrix elements (i.e., the 190 exchangeability values for each pair of amino acids) to sum to one for both analyses. Then we treated the normalized ***R*** matrix as a vector of 190 values and calculated a matrix of Euclidean distances among the models for the cluster analysis. We used three normalized vectors for the PCAs: 1) vectors with all 190 elements; 2) vectors of 75 elements limited to amino acid exchanges possible given single nucleotide change (1-nt interchanges); and 3) vectors of 101 elements limited to amino acid exchanges possible given two nucleotide changes (2-nt interchanges). We used JMP-Pro version 12.2 (SAS Institute Inc.) with default settings for the PCA.

We compared the 1-nt exchangeabilities for our clade-specific models to several matrices that describe amino acid properties. First, we used a symmetric version of the Yampolsky and Stoltzfus (2005) EX matrix, which describes the impact of mutations in laboratory mutagenesis studies. Note that the EX matrix is unrelated to the Le *et al*. (2008) EX2 model, which is an XB model (using our nomenclature). Lower EX matrix values indicate that mutating wild-type amino acid *i* to amino acid *j* typically results in more severe phenotypic changes in the laboratory. The EX_*s*_ matrix (EX matrix-symmetric) was produced by averaging the EX matrix values for *i* to *j* mutations and *j* and *i* mutations and then normalizing the matrix to assign the most experimentally exchangeable amino acid pair (I and V) a value of one. Second, we compared clade-specific model exchangeabilities to matrices that capture differences in amino acid side-chain volume and polarity, which were obtained from Braun (2018). All of these comparisons used Spearman’s rank correlations with two-tailed tests for significance.

## 3 Results and Discussion

### 3.1 Models trained on vertebrates and non-vertebrates form two clusters in model space

Clustering of Euclidean distances among models along with midpoint rooting revealed two distinct clusters in the model space (Fig. 2). The first cluster comprises the bird and mammal models and three standard empirical models trained using viral data. The second includes all of the other models estimated for this project along with all other standard empirical models. These results corroborate the hypothesis that patterns of sequence evolution vary across the tree of life and they further suggest that models trained using vertebrate data are especially distinctive.

**Fig. 2.**
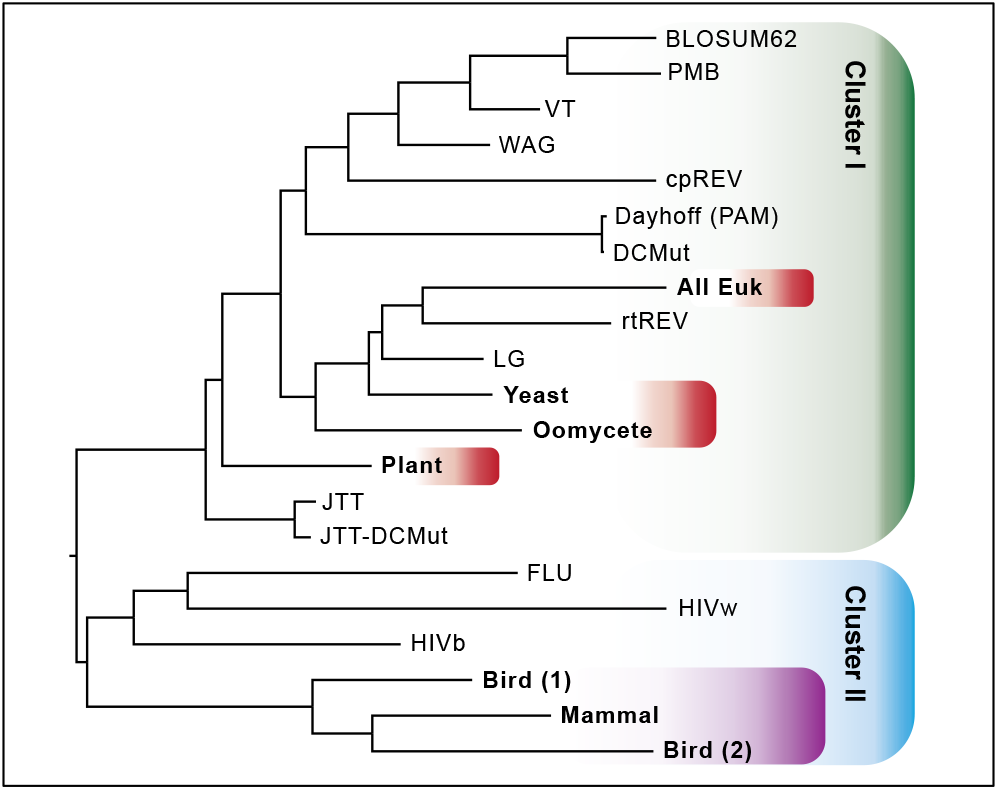
Cluster analysis for clade-specific models of sequence evolution. A ‘tree of models’ generated by neighbor-joining of Euclidean distances among exchangeability matrices for the new models (bold) and standard empirical models.

The strong separation between the vertebrate models and the other clade-specific models was also evident in a PCA of the 190 exchangeability parameters of these models (Fig. 3a). PC1 and PC2 were both significant, but PC1 explained most of the of variation and it separated the models into vertebrate and non-vertebrate models. Perhaps surprisingly, the three vertebrate models (two of which were estimated using bird data) appeared to be about as distinct as the non-vertebrate models. The PCA for the 1-nt exchangeability values (Fig. 3b) was quite similar to the all exchangeability PCA, probably reflecting the fact that the largest exchangeability values are those possible with a single substitution (Supplementary File 1). In contrast, PCA of 2-nt exchangeabilities (Fig. 3c) revealed a slightly different pattern; in that analysis PC1 also explained most of the variance but the plant model fell between the vertebrate models and the models for microbial eukaryotes (i.e., the yeast and oomycete). The vertebrate models were closer to each other than they were in the 1-nt PCA and the ‘all Euk’ model was located even further from vertebrates than the yeast and oomycete models. The latter finding probably reflects the fact that microbial eukaryotes dominate that dataset. The influence of 2-nt exchanges could explain the distances (sum of branch lengths between models) among vertebrate models and between vertebrate models and the plant and microbial eukaryote models in the cluster analysis (Fig. 2).

**Fig. 3.**
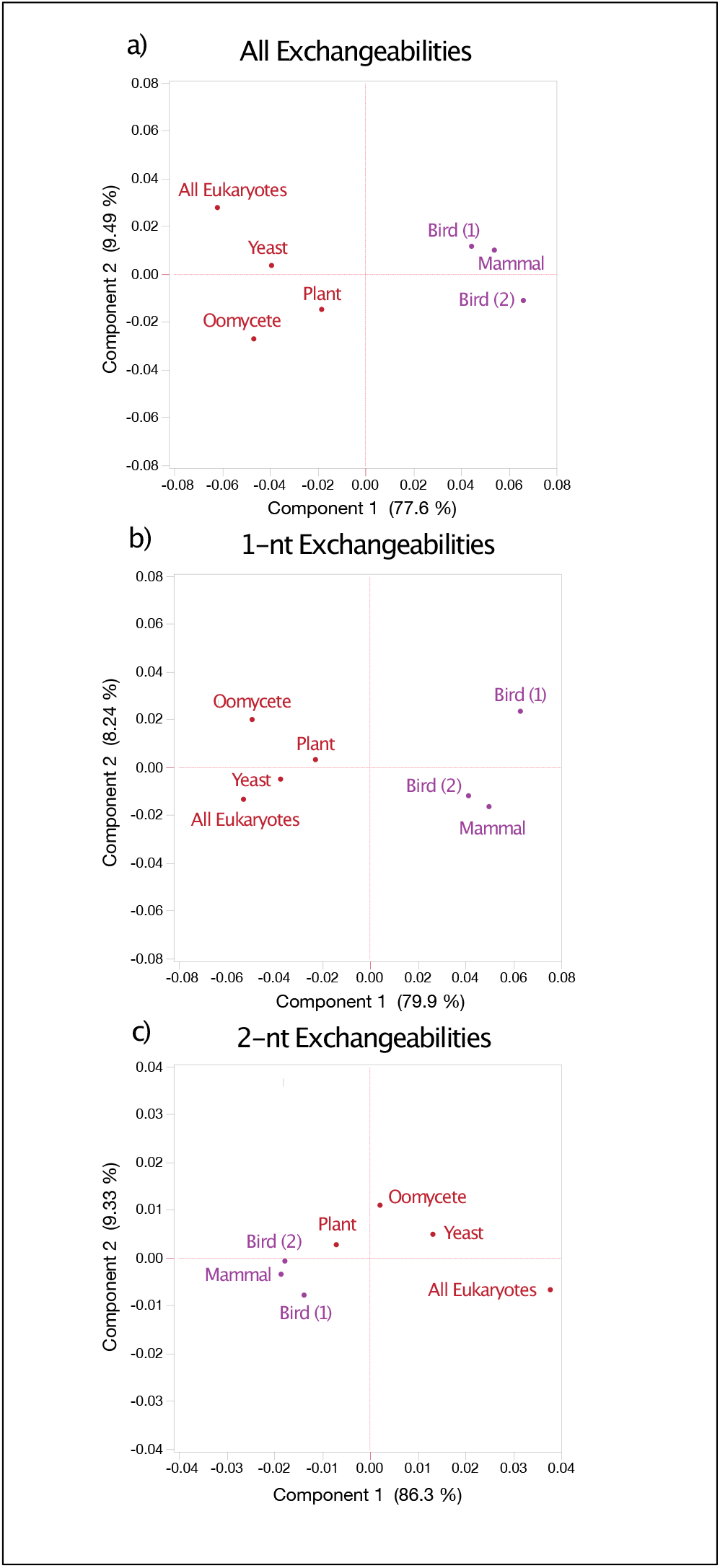
PCA of the clade-specific models of sequence evolution. Plot showing the first two PCs for analyses of exchangeability (***R*** matrix) parameter estimates. The PCAs were calculated using (a) all values; (b), 1-nt values; and (c) 2-nt values. The proportion of the variance explained by each PC is listed alongside each axis.

### 3.2 The best-fitting model for most individual proteins is the appropriate clade-specific model

The best-fitting model for individual proteins in each validation dataset was one of novel clade-specific models (Table 3); the only exception was the ‘all Euk’ validation dataset where the LG model (Le and Gascuel, 2008) had the best fit more than 60% of the time (compared to 31.5% for the new ‘all Euk’ model). The best-fitting models for the vertebrate validation datasets were split among the three vertebrate models (Table 3). The results for individual proteins in the training data were virtually identical (Supplementary Table S1); the exception was the ‘all Euk’ training data where the new ‘all Euk’ model had the best fit for 93.5% of proteins. These results indicate that the average patterns of protein evolution for each clade provide substantial information regarding the patterns of substitution within those clades and further suggests that idiosyncratic differences among proteins play a limited role in model fit (i.e., our results are consistent with the top portion of the model space shown in Fig. 1).

**Table 3.** Percentage of times clade-specific models were the best-fitting model for individual proteins in each validation dataset

| Best-fit model | Birds | Mammals | Plants | Oomycetes | Yeast | All Euk |
| --- | --- | --- | --- | --- | --- | --- |
| **Bird (1)** | **25** | 2 | 1 | — | — | — |
| **Bird (2)** | 24 | 21 | — | — | — | — |
| **Mammal** | 27.5 | **65** | — | — | — | — |
| **Plant** | 6.5 | 4 | **70.5** | 4.7 | 2.5 | — |
| **Oomycete** | 0.5 | — | 2 | **84** | 1.5 | 2 |
| **Yeast** | 1 | — | 3 | 2 | **88** | 4.7 |
| **All Euk** | — | — | 0.5 | 2 | 2 | **31.5** |
| LG | 1.5 | 0.5 | 5.5 | 5.3 | 4.5 | 60.4 |
| JTT | 8 | 6 | 16.5 | — | — | — |
| WAG | — | — | 0.5 | 1.3 | — | 1.3 |
| DCMut | — | — | 0.5 | — | — | — |
| BLOSUM | 0.5 | 0.5 | — | — | — | — |
| PMB | 1 | — | — | — | — | — |
| VT | 1.5 | — | — | — | — | — |
| HIVb | 2 | — | — | — | — | — |
| FLU | 0.5 | 0.5 | — | — | — | — |
| mt models | 0.5 | 0.5 | — | 0.7 | 1.5 | — |

### 3.3 Variation among structural environments is stronger than the variation among clades

Clustering the matrices from the exposed/buried model XB models with standard empirical models and the two matrices from the EX2 model (Le *et al*., 2008) revealed three relatively distinct clusters (Fig. 4). All exposed models formed a divergent cluster on one side of the midpoint root; the deepest split within the exposed models was between vertebrates and non-vertebrates (the EX2 exposed component nested within non-vertebrates). The results for the buried components was similar; the bird and mammal buried components formed a cluster that was distinct from the second group that included the buried components of all non-vertebrate models and the EX2 model. Both of those buried clusters were nested within groups of standard empirical models (none of the latter were structure aware). The vertebrate buried components formed a cluster sister to three viral models (HIVb, HIVw, and FLU). Thus, matrices for the structural models exhibited two levels of separation: 1) the separation between the exposed and buried clusters; and 2) the separation between the taxonomic groups (vertebrates vs. all other taxa).

**Fig. 4.**
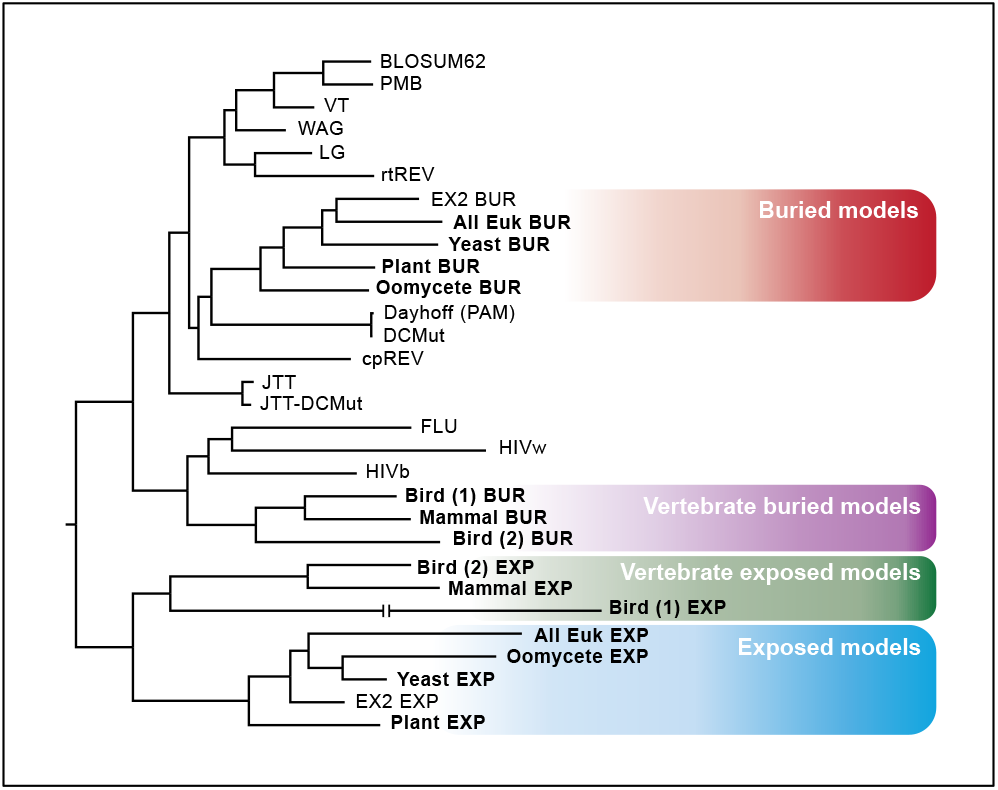
Cluster analysis for clade-specific models of sequence evolution that incorporate protein structure. Our structural models are mixture of two matrices, similar to the EX2 model of Le et al. (2008). This ‘tree of models’ was generated by neighbor-joining of Euclidean distances among exchangeability matrices for the new XB model components (bold), the two components of the EX2 model, and standard empirical models. The line break on the “bird (1) EXP” branch indicates that it was a long branch that was shortened for readability. This tree is available from github as a nexus format.

PCA of all 190 exchangeability parameters (Fig. 5a) and the 1-nt exchangeabilities (Fig. 5b) for the structural subsets of these datasets revealed similar patterns. In both cases PC1 explained ~50% of the variance and it separated the exposed and buried models. In contrast, PC2 explained slightly more than 25% of the variance and it separated the models by clade in a manner consistent with the models based on all data. The exposed ‘bird 1’ model, which was based on the protein alignments used in Jarvis et al. (2014), is especially distinctive. In contrast, PCA of amino acid exchangeabilities that require at least two nucleotide substitutions (2nt exchangeabilities) was less informative; most models clustered near the center of a score plot of the first two PCs (which together explain 82.2% of the variance among models, see Fig. 5c). The exposed and buried submodels of the ‘all Euk’ XB model were the most distinctive, with higher values of PC1 than any other sub-models from the same structural environment. The major similarity between the 2-nt PCA and the others is that the vertebrate exposed sub-models for vertebrates were more spread out than the buried sub-models for the same taxa, with the exposed ‘bird 1’ model based on the protein alignments used in Jarvis *et al*. (2014) being especially distinctive.

**Fig. 5.**
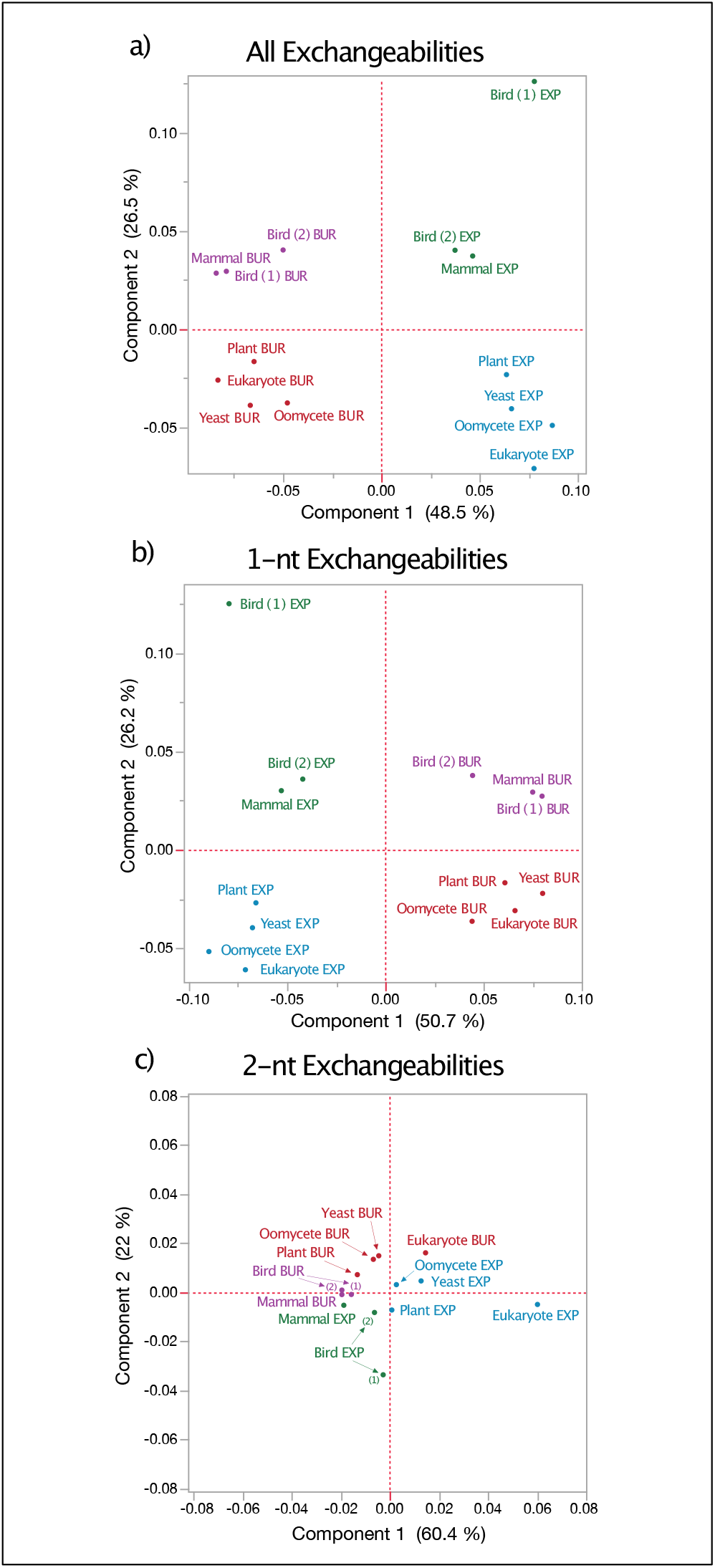
PCA of the clade-specific models of sequence evolution. Plot showing the first two PCs for analyses of exchangeability (***R*** matrix) parameter estimates. The PCAs were calculated using (a) all values; (b), 1-nt values; and (c) 2-nt values. The proportion of the variance explained by each PC is listed alongside each axis.

### 3.4 The best-fitting XB model for most individual proteins is the appropriate clade-specific XB model

The results validation set classification using clade-specific mixture models that were trained using exposed- and buried-site data as a classifier were similar to those obtained using the all-sites models (Table 4). The majority of the best-fitting XB model for the plant, oomycete, and yeast validation data were the appropriate clade-specific models (>75% in all cases). Likewise, the best-fitting models for proteins in the vertebrate validation sets were almost always models trained using vertebrate data (>80% in both case). As we observed with the all-sites models, the XB model was not our novel ‘all Euk’ XB model; it was the EX2 model (Le *et al*., 2008) instead. However, none of the truly clade-specific XB models were the best-fit to the ‘all Euk’ validation set.

**Table 4.** Percentage of times clade-specific XB models were the best-fitting model for individual proteins in each validation dataset

| Best-fit model | Birds | Mammals | Plants | Oomycetes | Yeast | All Euk |
| --- | --- | --- | --- | --- | --- | --- |
| **Bird (1)** | **26.5** | 5 | — | — | — | — |
| **Bird (2)** | 24.5 | 18 | — | — | — | — |
| **Mammal** | 32 | **68.5** | — | — | — | — |
| **Plant** | 12 | 8 | **94.5** | 4.7 | 1.5 | — |
| **Oomycete** | 1.5 | — | 1 | **76** | 5 | 6.7 |
| **Yeast** | 2.5 | — | 1 | 4 | **79** | 2 |
| **All Euk** | — | — | — | 3.3 | 1 | **33.5** |
| EX2 | 1 | 0.5 | 3 | 12 | 13.5 | 57.7 |

### 3.5 A ‘rule of opposites’ explains the exchangeabilities for different structural environments

The highest exchangeabilities for exposed sites involved pairs of hydrophobic residues; when exchangeabilities for all six models were averaged the three highest values corresponded to I-V, F-Y, and I-M. In contrast, the highest exchangeabilities for the buried environment were polar pairs (in this case, the three highest values were R-K, D-E, and Q-H). This pattern may seem surprising; after all, it has long been appreciated that polar residues are common in solvent exposed environments whereas hydrophobic residues dominate the buried sites (Worth *et al.*, 2009). We call the observation that the most exchangeable amino acids in each structural environment are the less common amino acids in that environment the ‘rule of opposites.’

The ‘rule of opposites’ allows us to differentiate between two alternative hypotheses to explain the relationship between exchangeabilities and amino acid frequencies. One might postulate that the amino acids that are rare in a specific structural environment would have very low exchangeabilities because those amino acid amino acids would be necessary for specific functions. Alternatively, one might postulate that exchanges between pairs of rare amino acids are common as long as the physicochemical nature of the amino acid is conserved. These results corroborate the second hypothesis and further suggest that at least some rare amino acids are especially exchangeable.

The exposed and buried sub-models of the XB models could be separated into a vertebrate and a non-vertebrate cluster (along PC2 in Fig. 5, panels a and b). Specific elements that separate the models were evident among the largest exchangeability values. Although the largest element for the exposed sub-models was I-V in both the vertebrate and plant/microbial groups, the next three elements differed. For vertebrates the next two elements involved exchanges between cysteine and aromatic residues (C-W and C-Y) whereas the plant/microbial models involved much more physicochemically-similar pairs (F-Y and L-M). Despite these differences, both groups conform to the rule of opposites (cysteine and aromatic residues are uncommon in solvent exposed environments; data available from github). In contrast, the top two exchangeabilities for the buried model were identical for the vertebrate and plant/microbial buried submodels, although there were certainly a number of additional differences.

### 3.6 Differences in the strength of selection appears to explain difference among clade-specific models

It should be possible to gain insights into the basis for the differences among-clade specific models by comparing changes in amino acid properties to the differences between vertebrate and plant/microbial models in their 1-nt exchangeabilities. Exchangeability differences for the all-sites models were correlated with experimental amino acid exchangeabilities (i.e., values in the EX_*s*_ matrix) and differences in side-chain volume. The correlation with EX_*s*_ was negative (Spearman’s correlation; *r*_*S*_ = −0.29309, *P* = 0.01071). The correlation with changes in amino acid side chain volume (hereafter, ∆ volume) was positive and even stronger (*r*_*S*_ = 0.39197, *P* = 0.00051). The directions of both correlations were consistent with the hypothesis that the major difference between vertebrate and non-vertebrate models is the relaxed selection against slightly deleterious mutations in vertebrates (presumably due to their lower long-term *N*_e_). In contrast, clade-specific model exchangeability differences were not correlated with ∆ polarity (*r*_*S*_ = −0.0405, *P* = 0.73009). These results suggest changes in side-chain volume is the primary property subject to differential selection between vertebrates and plants/microbial eukaryotes.

A similar pattern was evident for the exposed sub-model of the clade specific XB models. Specifically, surface residue exchangeability differences were correlated with EX_*s*_ (*r*_*S*_ = −0.34316, *P* = 0.00258) and ∆ volume (*r*_*S*_ = 0.45707, *P* = 4×10-5); those exchangeability differences were not correlated with ∆ polarity (*r*_*S*_ = −0.09805, *P* = 0.40265). The correlations were weaker for buried sites (*r*_*S*_ = −0.20979, *P* = 0.07085 for EX_*s*_; *r*_*S*_ = 0.30234, *P* = 0.00838 for ∆ volume; *r*_*S*_ = −0.03851, *P* = 0.7429 for ∆ polarity). However, comparisons of buried sub-models for vertebrate and plant/microbial did reveal some specific differences. 1-nt interchanges with higher buried-site exchangeabilities in vertebrates included A-T, R-H, M-V, R-Q, and C-Y whereas exchangeabilities with comparable elevation in the plant/microbial XB buried sub-models were F-Y, S-T, A-S, and N-H. Although there was no unifying physicochemical property for either set of pairs, we note that two of the pairs with elevated relative exchangeabilities in vertebrates (R-Q and C-Y) fall into different Dayhoff groups (i.e., the six groups shown in Fig. 84 of Dayhoff *et al*., 1978) and the pairs in the same Dayhoff group are relatively distinctive (e.g., R and H are both basic but they differ in shape, size, and even their charge at physiological pH). In contrast, two of the exchangeabilities elevated in the plant/microbial buried sub-models (F-Y and S-T) are physiochemically similar and only one (N-H) would change the Dayhoff group. Thus, these results are also consistent with the hypothesis that the lower long-term *N*_e_ of vertebrates has reduced the effectiveness of selection against slightly deleterious substitutions in vertebrates.

The hypothesis that among-clade differences in the patterns of protein sequence evolution reflects the strength of purifying selection raises several issues. First, the observation that our new vertebrate models clustered with models trained using viral data (HIVb, HIVw, and FLU; Nickle *et al.*, 2007; Dang *et al.*, 2010) seems puzzling if *N*_e_ is a major factor in establishing model differences; after all, viruses are microbes so one might assume their long-term *N*_e_ would be very large. However, the population biology of viral pathogens is complex, and drift appears to play an important role in their evolution (Kouyos *et al.*, 2006; Voronin *et al.*, 2009). Second, our failure to find a correlation between differences in exchangeabilities and ∆ polarity may seem surprising given the important role that polarity appears to play in models of protein evolution (cf. Braun, 2018). However, our goal was to examine differences between the exchangeabilities in vertebrate and plant/microbe models rather than the exchangeabilities themselves. An amino acid property that leads to similar exchangeability values in all taxa would not appear to be correlated with these differences. Thus, neither of those observations are problems for the hypothesis that differences in the strength of purifying selection can explain the differences among the clade-specific models.

### 3.7 Broadly sampled training data can distort model parameter estimates

Most empirical models have used as much training datasets as possible to reduce the variance of model parameter estimates. However, some studies have reported that estimates of parameters describing the amino acid substitution process exhibit time dependence (Benner *et al.*, 1994; Mitchison and Durbin, 1995; Müller *et al.*, 2002). The results of Benner *et al.* (1994), who estimated log-odds matrices using many pairs of aligned sequences selected to fall within certain divergence ranges, are especially interesting. They highlighted eight specific amino acid pairs; the log-odds scores for the first set (which we will call ‘type A pairs’) have higher values when they are estimated using divergent sequence pairs whereas the second set (hereafter, ‘type B pairs’) have lower log-odds scores they were estimated using divergent sequence pairs. Type A pairs (F-W, W-Y, C-M, and C-V) are similar amino acids (mean EX_*s*_ = 0.5258) that are encoded by codons that differ by at least two nucleotides. Type B pairs (C-W, R-C, C-Y, and R-W) are dissimilar amino acids (mean EX_*s*_ = 0.3547) encoded by codons that differ by a single nucleotide. These amino acid pairs led Benner *et al.* (1994) to conclude that “the genetic code influences accepted point mutations strongly at early stages of divergence, while the chemical properties of the side chains dominate at more advanced stages” (where ‘advanced stages’ refers to long evolutionary timescales).

We included the ‘all Euk’ training dataset to assess the impact of estimating model parameters using highly diverged sequences. Our exchangeability parameter estimates exhibited a pattern similar to the pattern observed by Benner *et al.* (1994) for log-odds scores; the mean exchangeability for type A pairs in the clade-specific models ranged from 15.4% of the ‘all Euk’ value (for W-Y) to 17.6% (for C-V). We observed similar patterns for both XB sub-models, with the mean exchangeabilities for type A pairs ranging from 13.2% of the ‘all Euk’ value (for buried site W-Y interchanges) to 22.1% (for exposed site W-Y interchanges). As expected, we observed the opposite pattern for type B pairs. When we normalized the ‘all Euk’ type B exchangeabilities to the maximum for that pair in any clade specific model we found values that ranged from an absolute minimum of 3.4% (for R-W interchanges in the exposed environment) to 24.8% (for R-C interchanges in the exposed environment).

Kosiol and Goldman (2011) pointed out that apparent time-dependence is problematic; after all, long-term substitution patterns ultimately reflect the accumulation of substitutions over many short periods of time. Thus, Benner *et al.* (1994) log-odds score estimates should not exhibit time dependence if the accumulation of amino acid substitutions can be modeled as a time-homogeneous Markov process. Kosiol and Goldman (2011) resolved this paradox by showing that a time-homogeneous Markov model for nucleotides can appear non-Markovian when the data are aggregated into the encoded amino acids. In fact, they even demonstrated apparent time-dependence of log-odds scores for the type A and B pairs qualitatively similar to the Benner *et al.* (1994) patterns (although there were only two pairs, C-M and C-V, that were similar in quantitative terms). Exchangeabilities for the LG model, which was trained using a taxonomically diverse dataset (Le and Gascuel, 2008), also exhibited the pattern of high values for type A pairs and (to a lesser degree) low values for type B pairs. Three of the type A pairs (F-W, W-Y, and C-M) in the LG model had values ~60% of the ‘all Euk’ comparable values and they were higher than the comparable values for any clade-specific model (see models on github). This suggests the LG model may be subject to a ‘time-dependency’ effect similar to our ‘all Euk’ model, albeit not as extreme.

The fact that type B pairs involve physicochemically-dissimilar amino acids that require a single nucleotide substitution for interchanges creates an additional complexity. They are exactly the type of substitutions expected to accumulate at an elevated rate in taxa where the long-term *N*_e_ is lower so the observation that vertebrate models always had the highest type B exchangeabilities (see models github) is not surprising. The surprise is actually the high values for the type A substitutions, which involve similar amino acids but require multiple substitutions. Some type B exchanges represent potential intermediates for type A substitutions (e.g., the only two-step pathway for F-Y involves C as an intermediate). Thus, one would expect strong selection against these disfavored intermediates to reduce type A exchangeabilities. However, type A exchangeabilities are quite high in the ‘all Euk’ model (e.g., W-Y was the highest 2-nt exchangeability, and it had the ninth highest value of the 190 exchangeabilities). Overall, the ‘all Euk’ models exhibit many similarities to the plant/microbial models (Figs. 2 and 4) combined with the distortion of some rate matrix parameter estimates (especially for type A pairs) superimposed. Those distortions of the rate matrix may have an impact on other uses of these rate matrices, like phylogenetic estimation.

## 4 Conclusions

Efforts to estimate models of protein sequence evolution began in the very earliest days of computational biology; the first version of the PAM matrix was estimated over 50 years ago using a mere 814 substitutions from 11 protein families (Dayhoff *et al.*, 1969). However, analyses of empirical models have provided little information about the processes governing protein evolution beyond the relatively straightforward conclusion that most amino acid exchanges involve physicochemically-similar amino acids. However, that was a conclusion that Dayhoff and Eck (1969) reached (in very general terms) by examining the first version of the PAM matrix. On the other hand, efforts to develop models of protein evolution from first principles (e.g., Parisi and Echave, 2001; Bastolla *et al.*, 2006; Arenas *et al*., 2013), remain impractical for phylogenetic analyses, especially in the phylogenomic era when hundreds or thousands of protein alignments are analyzed (e.g., the studies in Tables 1 and 2). The continued development of empirical models (e.g., Whelan and Goldman, 2001; Le and Gascuel, 2008) has provided models that can be used in that framework. It has not escaped our attention that our clade-specific models can also be used to improve phylogenomic analyses. The HIVb/HIVw (Nickle *et al*., 2007) and FLU (Dang *et al*., 2010) models were generated to improve analyses of proteins from those viruses; our clade-specific models should improve phylogenetic estimation for specific taxa. Moreover, our clade-specific XB models should further improve model fit (and tree estimation) by accommodating variation among taxa and variation among-sites within proteins due to protein structure. All of our models are available in from github and can be implemented in programs, such as IQ-TREE (Nguyen *et al*., 2015), that are used in many phylogenomic studies.

Although our models may be valuable for phylogenomic inference, the primary goal of this effort was to learn about the various ways that protein evolution has changed over time. Many efforts to understand the ways that evolutionary models change over time have assumed a single model for all sites within proteins. For practical reasons, they have also reduced the model dimension by focusing on single parameter, like the ratio of radical to conservative substitutions with radical vs. conservative substitutions in defined in a binary manner (e.g., Nabholz *et al*., 2013; Weber and Whelan, 2019). Although this basic approach has been extended to a limited number of parameters by considering the physicochemical properties of amino acids (Braun, 2018), it is difficult to ‘cast a wide net’ in order to learn the ways that the process of amino acid has changed over time. Herein, we have estimated parameters that describe protein evolution in various clades using a simple framework (the GTR_20_ model) that does not presuppose an important role for any specific amino acid property. In doing so we found that there is substantial variation among clades in their model and that this variation among clades is evident both for amino acids located on the surface of proteins and for residues buried in the interior of proteins. We also found evidence that vertebrates are more tolerant of substitutions that change amino side-chain volume than plant/microbial models; however, this was only evident in only for models that describe the evolution of solvent exposed residues. We also showed that training empirical models using sequences sampled from taxa that were sampled too broadly (i.e., the ‘all Euk’ training data) can lead to distorted parameter estimates. Finally, we found that most proteins from a specific taxon were clustered in model space and that a relatively the simple hypothesis – patterns of substitution reflect the strength of purifying selection, which differs among taxa due to differences among taxa in their long-term *N*_e_ – can explain many of the observed differences among taxa

## Acknowledgements

We are grateful to Rebecca Kimball, Gordon Burleigh, Gavin Naylor, Emily Sessa, and Tamer Kahveci for helpful conversations while this work was ongoing.

## Supplementary data

**Table S1.** Percentage of times clade-specific models were the best-fitting model for individual proteins in each training dataset

| Best-fit model | Birds (1) | Birds (2) | Mammals | Plants | Oomycetes | Yeast | All Euk |
| --- | --- | --- | --- | --- | --- | --- | --- |
| **Bird (1)** | **49.2** | 4.4 | 5.2 | 0.3 | — | — | — |
| **Bird (2)** | 16.8 | **72** | 23.4 | 0.3 | — | — | — |
| **Mammal** | 14 | 14 | **59.1** | 0.3 | — | — | — |
| **Plant** | 9.2 | 3.6 | 5.7 | **88.1** | 1.1 | 1 | — |
| **Oomycete** | 0.8 | 0.8 | 0.8 | 5.5 | **86.5** | — | 0.4 |
| **Yeast** | — | 0.4 | — | 2.6 | 1.1 | **96.5** | — |
| **All Euk** | — | — | — | 0.6 | 4 | 1 | **93.5** |
| LG | — | 0.4 | — | 0.6 | 3.6 | 1 | 5.2 |
| JTT | 3.2 | 1.6 | 4 | — | 0.4 | — | — |
| JTT-DCMut | 0.4 | — | 1.2 | 1.3 | 0.4 | — | — |
| WAG | 0.4 | — | — | 0.6 | 0.7 | — | 0.4 |
| Dayhoff | — | 0.4 | — | — | — | — | — |
| BLOSUM | 3.2 | — | — | — | 0.4 | — | — |
| VT | 1.6 | — | — | — | 0.4 | — | — |
| HIVb | 0.4 | 0.4 | — | — | — | — | — |
| FLU | 0.4 | 2 | — | — | — | — | — |
| rtREV | — | — | — | — | 0.4 | — | — |
| mt models | 0.4 | — | 0.4 | — | 1.1 | 0.5 | 0.4 |

Our clade-specific models are presented in bold. Values are rounded to the nearest 0.1%; ‘—’ indicates models that never had the best fit to any protein the specified validation dataset. Cases where mitochondrial models had the best fit were grouped (‘mt models’). This table only reports the best-fitting rate matrix; each protein was allowed to have any among-sites rate heterogeneity model.

**Table S2.**
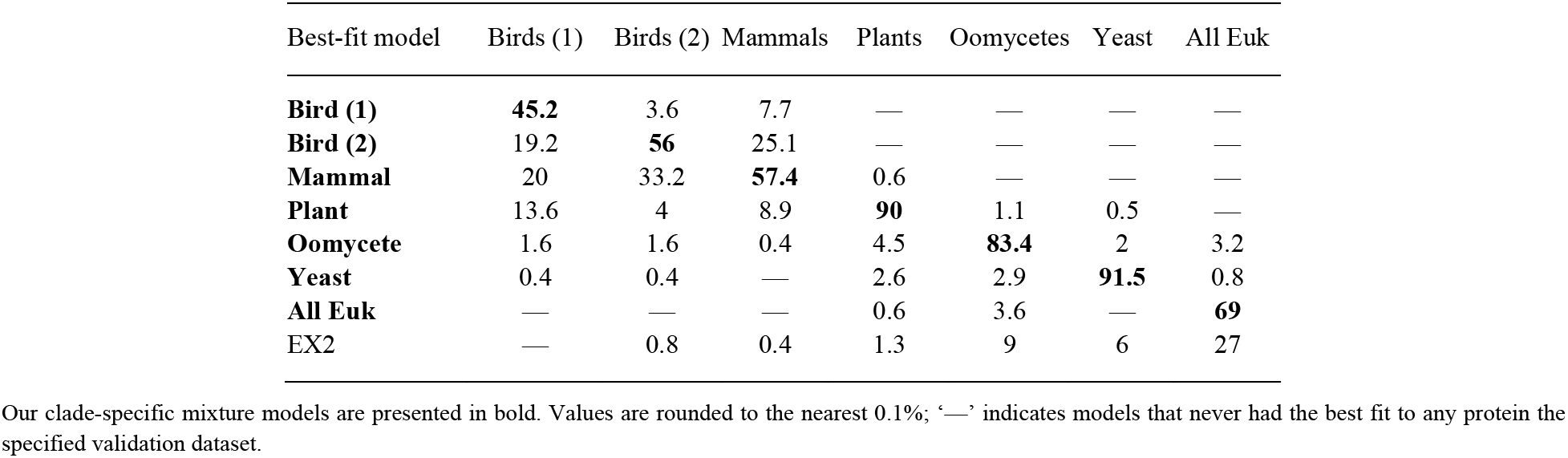
Percentage of times clade-specific XB mixture models were the best-fitting model for individual proteins in each training dataset

| Best-fit model | Birds (1) | Birds (2) | Mammals | Plants | Oomycetes | Yeast | All Euk |
| --- | --- | --- | --- | --- | --- | --- | --- |
| **Bird (1)** | **45.2** | 3.6 | 7.7 | — | — | — | — |
| **Bird (2)** | 19.2 | **56** | 25.1 | — | — | — | — |
| **Mammal** | 20 | 33.2 | **57.4** | 0.6 | — | — | — |
| **Plant** | 13.6 | 4 | 8.9 | **90** | 1.1 | 0.5 | — |
| **Oomycete** | 1.6 | 1.6 | 0.4 | 4.5 | **83.4** | 2 | 3.2 |
| **Yeast** | 0.4 | 0.4 | — | 2.6 | 2.9 | **91.5** | 0.8 |
| **All Euk** | — | — | — | 0.6 | 3.6 | — | **69** |
| EX2 | — | 0.8 | 0.4 | 1.3 | 9 | 6 | 27 |

Our clade-specific mixture models are presented in bold. Values are rounded to the nearest 0.1%; ‘—’ indicates models that never had the best fit to any protein the specified validation dataset.

## Additional files

Model files and other information are available from https://github.com/ebraun68/clade_specific_prot_models

